# Selective oxytocin receptor activation prevents prefrontal circuit dysfunction and social behavioral alterations in response to chronic prefrontal cortex activation in rats

**DOI:** 10.1101/2022.11.08.515590

**Authors:** Philipp Janz, Frederic Knoflach, Konrad Bleicher, Sara Belli, Barbara Biemans, Patrick Schnider, Martin Ebeling, Christophe Grundschober, Madhurima Benekareddy

## Abstract

Social behavioral changes are a hallmark of several neurodevelopmental and neuropsychiatric conditions, nevertheless the underlying neural substrates of such dysfunction remain poorly understood. Building evidence points to the prefrontal cortex (PFC) as one of the key brain regions that orchestrates social behavior. We used this concept with the aim to develop a translational rat model of social-circuit dysfunction, the chronic PFC activation model (CPA). Chemogenetic designer receptor hM3Dq was used to induce chronic activation of the PFC over 10 days, and the behavioral and electrophysiological signatures of prolonged PFC hyperactivity were evaluated. To test the sensitivity of this model to pharmacological interventions on longer timescales, and validate its translational potential, the rats were treated with our novel highly selective oxytocin receptor (OXTR) agonist RO6958375, which has a significantly improved pharmacokinetic profile over oxytocin. CPA rats showed reduced sociability in the three-chamber sociability test, and a concomitant decrease in neuronal excitability and synaptic transmission within the PFC as measured by electrophysiological recordings in acute slice preparation. Sub-chronic treatment with a low dose of the novel OXTR agonist following CPA interferes with the emergence of PFC circuit dysfunction, abnormal social behavior and specific transcriptomic changes. These results demonstrate that sustained PFC hyperactivity modifies circuit characteristics and social behaviors in ways that can be modulated by selective OXTR activation and that this model may be used to understand the circuit recruitment of prosocial therapies in drug discovery.

## Introduction

Social impairment is a transdiagnostic hallmark of different neurodevelopmental and neuropsychiatric conditions such as autism spectrum disorders (ASD) and schizophrenia (SZ). In a healthy brain, social behavior is tightly controlled by the PFC, which integrates sensory information to create mental representations that influence social decision-making and actions (reviewed in (1,2)). Given its intricate connectivity not only with cortical regions involved in sensory processing and motor control, but also with subcortical brain areas of the limbic system (3,4), such as the nucleus accumbens, ventral tegmental area (VTA), lateral habenula (LHb) or the amygdala, the PFC is ideally placed to elicit top-down influence on attention, motivation and affect, which is highly-relevant for social behavior. In ASD and SZ, converging evidence from neuroimaging, physiological and systems neuroscience point to PFC dysfunction (5–7) that may represent a common substrate for disease-related social deficits. On the mechanistic level, the disbalance of PFC excitation-to-inhibition has been extensively studied and shaped our current understanding of how complex genetics and environmental factors contribute to social deficits in psychiatric disorders (8–10). Particularly the impairment of cortical interneurons appears as a central feature in both SZ (11) and ASD (12). In fact, several mechanistic rodent studies impressively showed that shifting excitation-to-inhibition balance in the PFC e.g. by silencing interneurons (13) or by exciting pyramidal neurons (14) disrupts socially-directed behavior.

From a therapeutic perspective, several pharmacological approaches are being explored that target social brain circuits, with the hope to alleviate disease-associated social deficits. One of these approaches relies on the application of oxytocin, a neuropeptide that is particularly well-known for its role in modulating social behavior (reviewed in (15–17)) and anxiety (18). However, despite promising preclinical results (19–23), intranasal application of oxytocin in autistic children showed no improvement of social and cognitive function (24), demonstrating the need to better understand how oxytocin affects brain circuits governing social behavior. In the mammalian brain the major sources of oxytocin are hypothalamic neurons projecting to widely distributed brain areas, including the PFC (25). Oxytocin that is released upon social cues may, beside its well-known neuroendocrine effects (26), have a wide-spread influence on brain networks. In the PFC, oxytocin activates interneurons that, in turn, inhibit either layer 2/3 pyramidal neurons in male mice to reduce anxiety, or layer 5 pyramidal neurons in female mice to elicit prosocial behavior (27). For the hippocampus it has been shown that oxytocin enhances signal transmission by increasing the activity of fast-spiking interneurons (28). Similarly, oxytocin boosts inhibition in the amygdala to attenuate fear responses (25). In the VTA oxytocin increases action potential firing of nucleus accumbens-projecting dopaminergic neurons to gate social reward (29). Treatment with oxytocin has been shown to rescue social deficits in a variety of rodent models relevant for ASD (19–23). However, from these studies it remains unclear if oxytocin can restore social behavior by acting specifically on OXTR in prefrontal circuits given its short half-life and its pharmacological effect on the vasopressin1a receptor (V1aR).

Therefore, in our study we developed a PFC circuit model for chronic social impairment, explored the underlying cellular mechanisms and tested its utility for drug development. In this context, we describe for the first time a selective, brain-penetrant peptidic OXTR agonist that was capable of preventing social-circuit dysfunction observed in our model.

## Methods

### Animals

Adult Sprague Dawley rats (Charles River, Saint-Germain-sur-l’Arbresle, FR) were maintained on a reversed 12-hour light/dark cycle with food and water ad libitum. Experiments were conducted in adherence to the Swiss federal ordinance on animal protection.

### Viral injections

Rats were deeply anesthetized and hM3Dq-expressing Adeno-associated viruses (AAV) packaged by the Viral Vector Facility of the ETH Zürich were injected into the medial PFC. For control rats, a control virus carrying only the genomic sequence for the fluorophore was used. In a subset of rats a retrograde-infecting EGFP-expressing AAV was additionally injected in the dorsomedial thalamus to label PFC projection neurons. Details are available in the supplementary material.

### Peptide synthesis

RO6958375 was synthesized via microwave-based solid phase Fmoc-chemistry using the Liberty Lite system (CEM, Matthews, US). The synthesis was carried out using TentalGel-S RAM resin as a solid support, involving alloc- and allyl-cleavage. For on-bead cyclisation, the coupling-reagent was added to the resin. Completion of cyclisation was verified via ninhydrin test. After cleaving the peptide from the resin, the cleaved peptide was precipitated from cold ether, centrifuged and dissolved in water/acetonitrile and lyophilized. Finally, the crude peptide was purified by HPLC and analyzed via Electrospray Mass Spectrometry. Details are available in the supplementary material.

### Pharmacological characterization of OXTR agonist RO6958375

RO6958375 was profiled in a calcium flux *in vitro* assay on cells expressing human or rat OXTR or the related human or rat vasopressin V1a, V1b and V2 receptors as previously described (30). Details are available in the supplementary material.

### Peptide *in vivo* distribution studies

Rats were administered RO6958375 subcutaneously at 0.03, 0.1, 0.3, or 4 mg/kg. Serial blood and one CSF sample were collected from each animal under deep isoflurane-anesthesia. Blood samples were collected from 0.25 to 4.5 h post-dose, and CSF sampling from 0.8 to 4.5 h. Compound concentrations in blood plasma and CSF were determined by means of LC–MS/MS. Details are available in the supplementary material.

### Pharmacological treatment

Two weeks after surgery, the hM3Dq ligand clozapine-*N*-oxide (CNO; BML-NS105, Enzo Life Sciences, Farmingdale, USA) was injected at a dose of 1 mg/kg s.c. on a daily basis for 10 days, followed by a wash-out phase of 7 days. For RO6958375 experiment CNO was provided in drinking water with a calculated final dose of 2 mg/kg. To test the effect of pharmacological intervention, the peptidic OXTR agonist RO6958375 (or a vehicle control) was injected daily during the CNO wash-out phase. The dose was 0.07 mg/kg (s.c.) if not noted otherwise.

### Three-chamber sociability test

Social behavior was assessed in a 3-chamber task from Noldus Ethovision (Noldus, Wageningen, NL) using established protocols reported previously (14). Details are available in the supplementary material.

### Electrophysiology

Acute slice electrophysiology was performed on 350 μm thick frontal sections using a VT1000S vibratome (Leica, Wetzlar, GER). Signals were recorded with a MultiClamp 700B and pClamp software (Axon Instruments, Molecular Devices, San Jose, California, US).

For testing synaptic transmission a bipolar stimulation electrode was located in layer 2/3 and postsynaptic field potential (fPSP) were recorded at 150 μm depth in layer 5. Stimulation currents were generated with an STG3000 and MC Stimulus II software (Multi Channel Systems, Reutlingen, Germany). Several protocols were performed: i.) Input-output protocol applying incremental current steps until 300 pA, ii.) paired pulse protocol using different interval durations and iii.) a synaptic short term depression protocol by delivering pulses of various frequencies. Presynaptic fiber volley (FV) and the fPSP were semi-automatically quantified using pClamp software and averaged per animal (three slices per animal, three trials for each slice).

Spontaneous activity was recorded in the central LHb and the VTA. After an active neuron was identified and waiting 5 min for the spike rate to normalize, action potential firing was recorded for at least 10 min. The firing frequency was analyzed for each neuron by using an adjustable threshold for spike detection in pClamp.

For patch-clamp experiments PFC layer 5 pyramidal neurons were recorded in whole-cell mode. After establishing a gigaseal, the membrane was opened and protocols were started after 10 min, to allow for sufficient exchange of intracellular solution and normalization of cell physiology. The membrane potential was held at −70 mV, and only neurons for which leak currents did not exceed ±100 pA and for which the axial resistance did not show changes higher than 20% were included for analysis. Here, we assessed i.) passive membrane properties ii.) steady-state potassium currents, and iii.) Intrinsic excitability applying step-wise current injections. Details are available in the supplementary material.

### RNA Sequencing

The mPFC was dissected out from the sections after electrophysiology, and the tissue was flash frozen. In brief, RNA was extracted, the quality was checked and concentration determined. 500 ng of RNA was used as input for library preparation using the Illumina TruSeq (Illumina, San Diego, US). Libraries were quality checked, pooled and sequenced on a HiSeq4000 (Illumina, San Diego, US). Details are available in the supplementary material.

### Statistical analysis

Behavioral and electrophysiological data was tested statistically with Prism 7 software (GraphPad Software Inc., La Jolla, USA). Applied statistical tests are described in the figure captions. Numeric values and statistical tests for key results are provided in supplementary table 1. Transcriptome data was analyzed using one-way ANOVA followed by a post-hoc test corrected for multiple comparisons using the original FDR method (31).

## Results

### CPA impairs socially-directed behavior

To increase the neuronal activity of the prefrontal network over many days, hM3Dq was expressed in PFC pyramidal neurons, and repeatedly activated over 10 days (Fig. 1A). Social behavior was assessed at baseline and one week after the phase of prefrontal hyperactivity using the three-chamber sociability task (Fig. 1B). We found that CPA rats, whose PFC was repeatedly activated, showed a strong trend toward fewer interactions with an unfamiliar conspecific (Fig. 1C). Control rats that were transduced only with the control fluorophore, exhibited no group level decrease in social interaction. Comparison between cohorts revealed that the sociability index is significantly decreased following CPA compared to controls (Fig. 1C).

**Figure 1:**
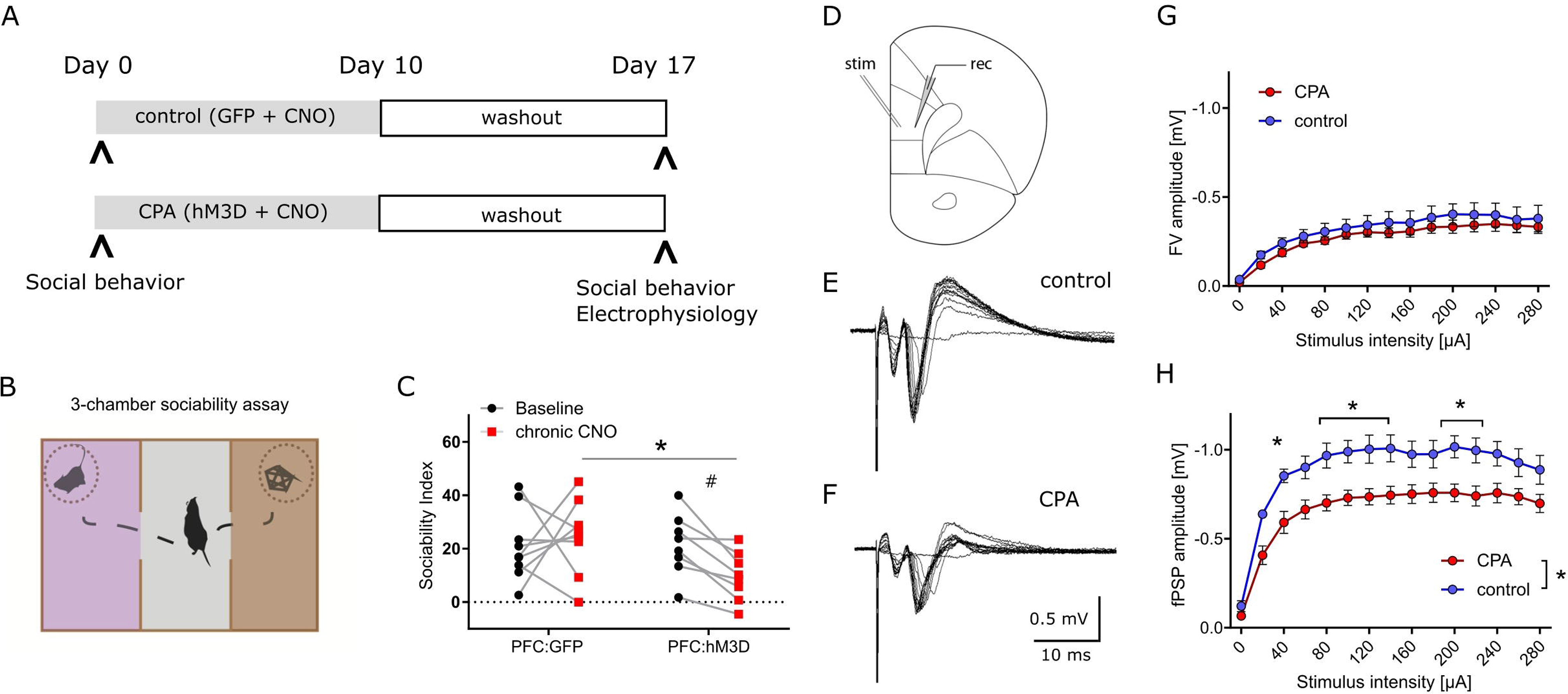
Chronic prefrontal activation impairs social behavior and PFC synaptic transmission. A) Schematic of experiment to induce prolonged activation of the prefrontal cortex using a combination of hM3Dq expression in the excitatory neurons and repeated administration of the ligand CNO. B) Cartoon representation of the 3-chamber sociability task used to measure social preferences. C) Sociability index measurements after chronic PFC activation. The control group was transduced with AAV:GFP while the treatment group was transduced with AAV:hM3Dq. Social behavior was measured before and after chronic CNO treatment. ^#^ p = 0.066 using two-way ANOVA followed by Fisher’s LSD test. Group comparison was done using Sidak’s post-hoc test * < 0.05. D) Schematic of experimental design: Extracellular stimulation was performed in PFC layer 2/3 and evoked responses were recorded in PFC layer 5 of acute slice preparations. E, F) Representative evoked responses under control and chronic prefrontal activation (CPA) conditions. Note that for individual current steps (0-280 uA) resulting voltage traces are overlaid. G, H) Quantitative analysis of two components of evoked responses (fiber volley, FV; postsynaptic field potential, fPSP) reveals a significant reduction in fPSP amplitudes for the CPA group. Data is displayed as mean ± SEM. Two-way ANOVA with Sidak’s post-hoc test was performed for statistical comparison. To compare the overall difference between control and CPA conditions, respective AUCs were tested with Student’s t-test. * p < 0.05.

### Prefrontal synaptic transmission is disrupted following CPA

Having revealed a chronic impairment of social behavior upon CPA, we aimed to elucidate the underlying cellular and circuit changes. First, we investigated synaptic transmission within the PFC in acute slices in which cells in layer 2/3 were stimulated with a bipolar electrode, while evoked potentials were recorded in layer 5 (Fig. 1D). Remarkably, the amplitudes of evoked field potentials were significantly decreased in CPA rats compared to controls (Fig. 1H). Fiber volley amplitudes showed a similar trend (Fig. 1G). To assess whether the decrease in neuronal responses might be a result of changed inhibitory feedback or presynaptic release properties, we also performed paired-pulse experiments and short-term depression experiments (Supplementary fig. 1). Neither the paired-pulse ratios nor synaptic depression were different between the two experimental groups, showing that local inhibitory circuits and presynaptic plasticity mechanisms remain intact following CPA.

#### CPA decreases excitability of prefrontal projection neurons

Next, we addressed putative changes on the single-cell level that could explain the impaired synaptic transmission by performing whole-cell patch-clamp recording of layer 5 pyramidal neurons (Fig. 2). To be able to record specifically from PFC projection neurons, we labeled these cells by infusion of a retrograde-traveling reporter virus targeting the mediodorsal thalamus and habenula (Fig. 2A). PFC projection neurons from CPA rats showed a significantly hyperpolarized resting membrane potential and reduced membrane resistance (Fig. 2B, C), as well as reduced intrinsic excitability upon current injections with a nearly doubling of the rheobase (Fig. 3H-I). In line with the reduced excitability, we found that subthreshold current injections in neurons from CPA rats resulted in smaller increases of the membrane voltage and decreased input resistance (Fig. 2J-N). Since the membrane potential and input resistance is critically regulated by several types of potassium channels, we inferred steady-state potassium currents. In particular, voltage-gated potassium currents elicited at −40 mV were strongly elevated and, as expected, the magnitude of these currents positively correlated with the rheobase (Supplementary fig. 2C, D). Similar differences were also evident for potassium currents elicited with 0 mV holding potential (Supplementary fig. 2G, H).

**Figure 2:**
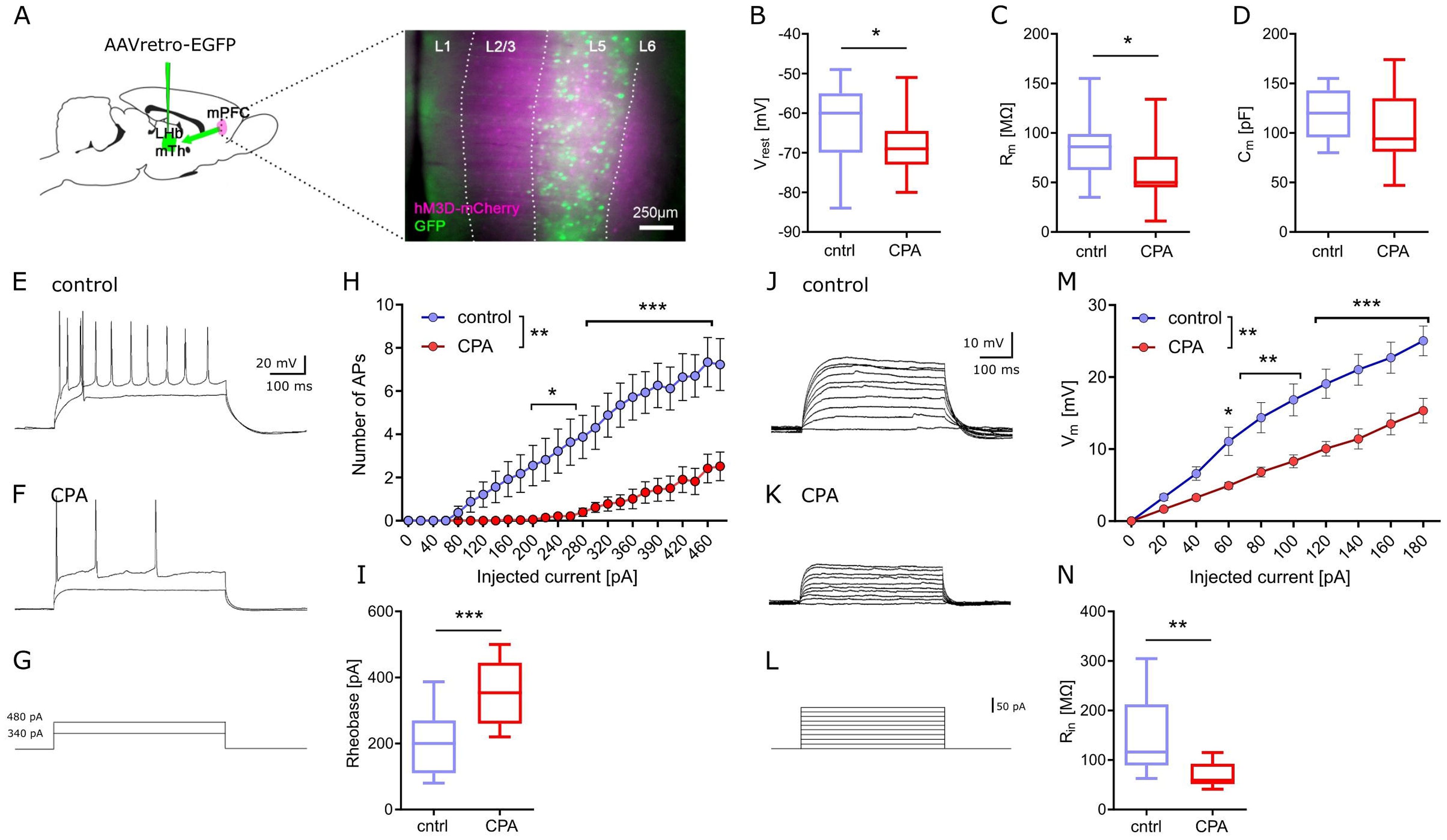
Chronic prefrontal activation reduces excitability of PFC output neurons. A) Schematic for whole-cell recording of PFC output neurons: Infusion of a retro-AAV reporter in the region of the lateral habenula (LHb) and the mediodorsal thalamus (MTh) allowed to perform whole-cell recordings from retrogradely-labeled PFC projection neurons (green) also expressing hM3Dq-mCherry (magenta) . B-D) Quantitative analysis of membrane properties (resting membrane potential, Vrest; membrane resistance, Rm; membrane capacitance, Cm) indicate that PFC projection neurons are more hyperpolarized and show a lower membrane resistance following chronic hyperactivation. To test whether these intrinsic changes affect neuronal excitability, current injections were performed. E-G) Representative voltage traces for 340 and 480 pA current steps. H) Quantitative analysis of the number of resulting action potentials (APs) reveals that the intrinsic excitability is dramatically reduced under CPA conditions, resulting in I) an increase in the rheobase. J-L) Representative traces for subthreshold voltage steps and resulting membrane potential, and M) corresponding quantification showing the decreased Input-output relation following CPA. N) Input resistance calculated for 80 pA. Students’s t-test was performed to test for statistical differences between groups. Two-way ANOVA with Sidak’s post-hoc test was performed for statistical comparison in H) and M). * < 0.05, ** < 0.01, *** < 0.001. Data in box-and-whiskers plots is displayed as the median with min. to max. All other data is displayed as mean ± SEM (box).

**Figure 3:**
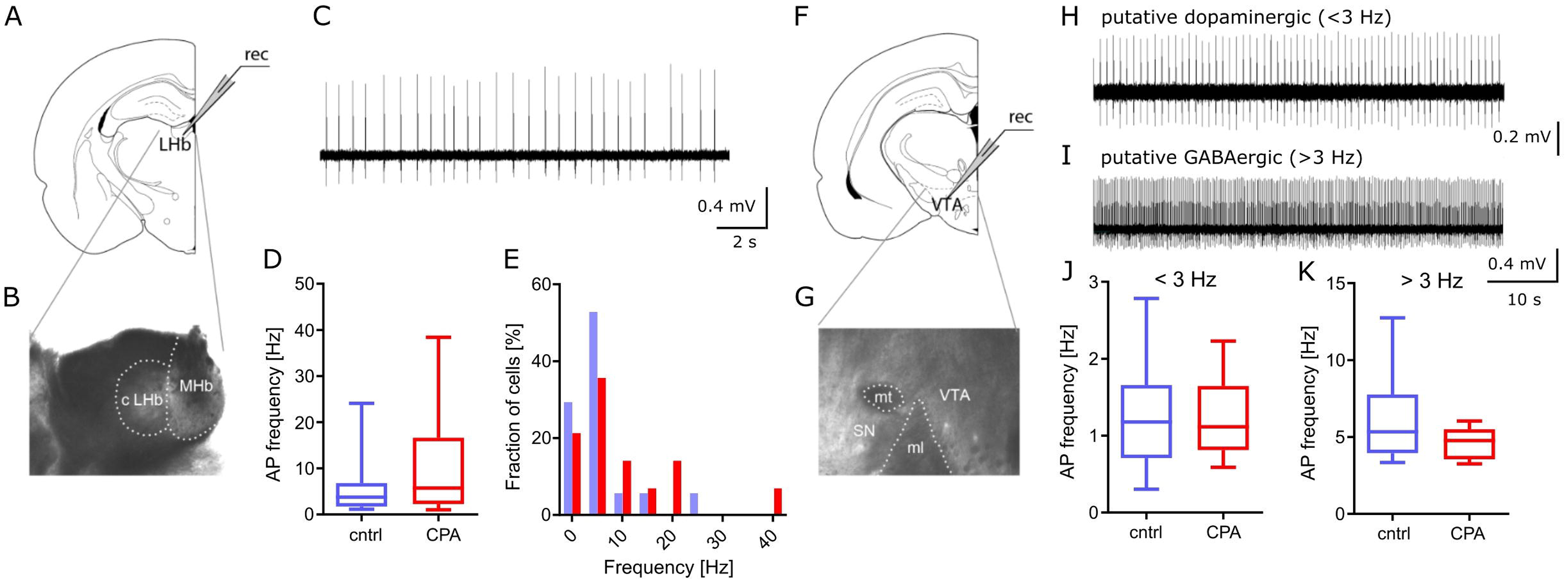
Spontaneous activity in PFC downstream targets LHb and VTA remains unchanged. A, B) Schematic and representative microscopy image of the location for extracellular recording in the LHb. C) Representative voltage trace from a spontaneously active neuron. D, E) Respective quantitative analysis of the AP frequency and the distribution of spontaneously active cells, shows no significant difference between the control and CPA group. F, G) Schematic and representative microscopy image of the location for extracellular recording in the VTA. H, I) Representative voltage traces from a putative dopaminergic cell (slow spiking, < 3 Hz) and a putative GABAergic (> 3 Hz spiking) neuron. J-K) Quantitative analysis of AP frequencies for both cell populations shows no significant groups differences. Statistical comparison was performed with Student’s t-test. Data is displayed as the median with min. to max. (box-and-whiskers).

### Spontaneous activity of PFC-downstream targets LHb and VTA remains intact upon CPA

Going beyond the PFC, we studied the function of two key downstream targets relevant in social behavior, the LHb and the VTA (Fig. 3). Considering that a large fraction of neurons within the LHb and VTA are spontaneously active, we performed extracellular recording of action potentials. Our results show that CPA does not lead to long-term changes of firing rates, in the LHb or the VTA (Fig. 3D-K), suggesting that these PFC-downstream targets may not be affected by CPA.

### Pharmacological and pharmacokinetic characterization of the novel peptidic OXTR agonist RO6958375

To assess the therapeutic potential of OXTR activation on social circuit impairment following CPA, we tested our novel selective OXTR agonist RO6958375. An extensive structure-activity relationship study of oxytocin revealed that substitution of the cystine bridge by a lactam (Fig. 4A) resulted in an unprecedented selectivity against vasopressin receptor 1a (V1aR). *In vitro* functional studies demonstrated that RO6958375 is a full OXTR agonist and equipotent to oxytocin on rat OXTR (Fig. 4B), while it is inactive on rat V1aR up to 1 μM. In contrast, oxytocin fully activated rat V1aR at 1 μM compared to the endogenous full agonist vasopressin (Fig. 4C). RO6958375 also fully activated the human OXTR and was inactive on human V1aR up to 27 μM (data not shown). EC_50_ values are provided in supplementary table 2. Furthermore, RO6958375 did not show any significant radioligand displacement or functional inhibition in a panel of 45 human receptors, channels, transporter and enzymes (see supplementary table 3).

**Figure 4:**
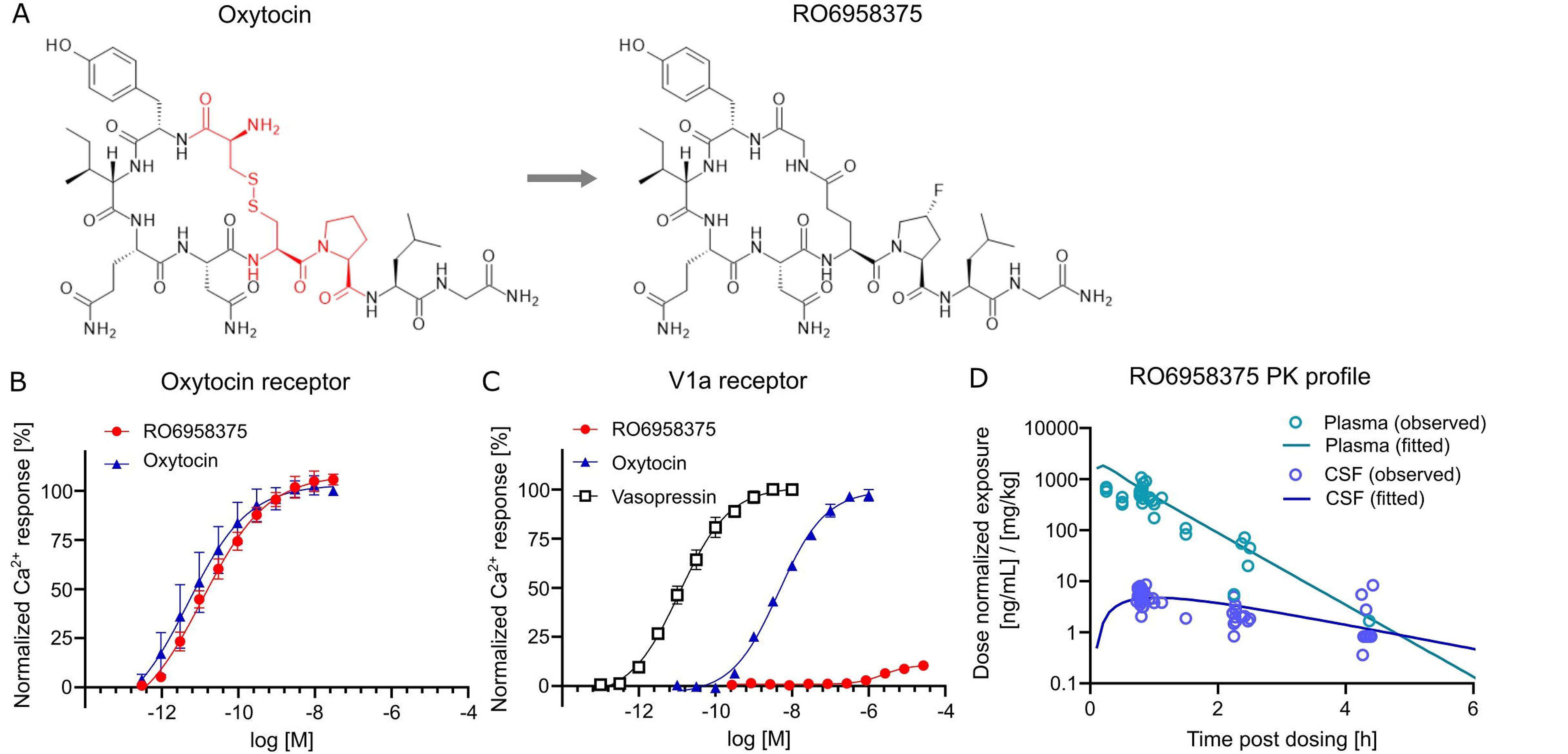
OXTR activation and pharmacokinetics of RO6958375. A) Chemical structure of oxytocin (with the changed amino acids highlighted in red) and our novel selective OXTR agonist RO6958375. B-C) Activity data of oxytocin and RO6958375 in rat OXTR and V1aR expression systems. D) *In vivo* SC pharmacokinetics of RO6958375 in rat plasma (negligible plasma protein binding) and CSF from different studies. Data is displayed as mean ± SEM.

For *in vivo* distribution studies the CNS uptake of RO6958375 after subcutaneous (SC) administration was tested in various single-dose pharmacokinetic studies in rats. Subcutaneous absorption was rapid, with a half-life in plasma of ca. 10-15 min. Interestingly, the CSF kinetic results indicated a longer residence of RO6958375 in brain (half-life of ca. 70 min) with a CNS uptake of ca. 1% from plasma (Fig. 4D). From the above studies, the highest concentrations detected in rat CSF at ca. 0.8 h after single SC administration of 0.03 or 0.1 mg/kg was equivalent to 0.24 and 0.58 nM, which is more than 10-fold above the *in vitro* OXTR EC_50_. RO6958375 is therefore sufficient to fully activate the rat OXTR for 4.3 h post-dose. For *in vivo* studies we selected subcutaneous doses of 2.4*10E-8, 0.001 and 0.07 mg/kg, leading to 0, 10 or 72% OXTR activation respectively, based on the *in vitro* OXTR dose-response.

### Sub-chronic selective OXTR activation prevents CPA-induced prefrontal circuit dysfunction

First, we tested whether OXTR activation can prevent the PFC dysfunction following CPA. Therefore, rats were repeatedly-injected with our novel selective peptidic OXTR agonist RO6958375 (RO, 0.07 mg/kg) or the vehicle for seven days, starting directly after CPA (Fig. 5A). Subsequent to the treatment, whole-cell recording of PFC layer 5 pyramidal neurons was performed (one rat per day) and tissue was preserved for later transcriptomic analysis. Remarkably, we found that the clear differences in passive membrane properties, evident between control rats and CPA rats, are not observed in RO6958375-treated CPA rats (Fig. 5C-E). Again, we also probed the intrinsic excitability by applying incremental current injections and found that RO6958375 treatment also fully restored the rheobase (Fig. 5K), whereas the number of evoked action potentials is only partially recovered for lower current steps (Fig. 5J). Consistent with the RO6958375-mediated restoration of membrane properties and intrinsic excitability, we also found that both, the current-membrane voltage relationship and the input resistance were normalized (Fig. 5L-M).

**Figure 5:**
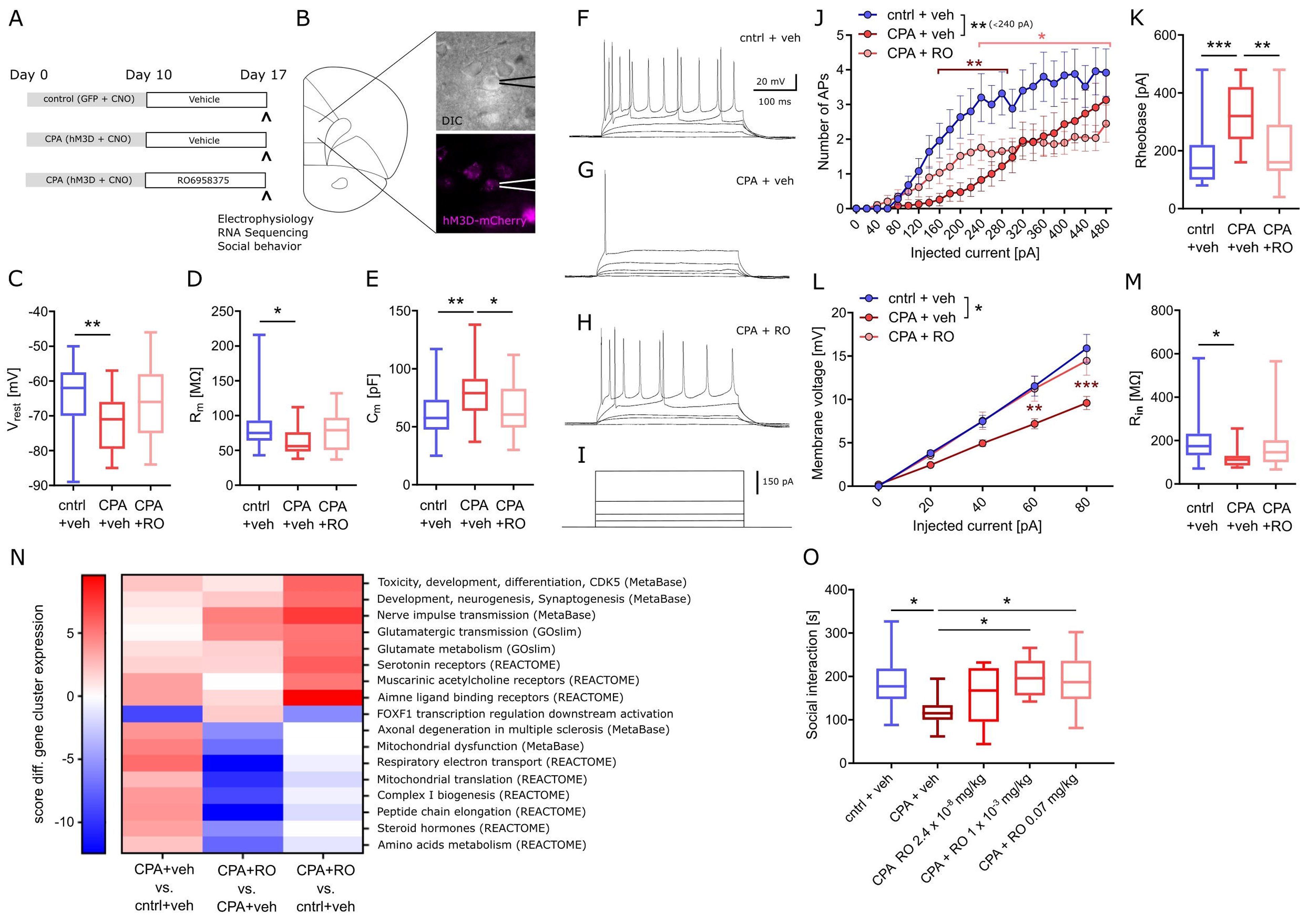
Chronic OXTR agonist treatment prevents circuit dysfunction following PFC overactivation. A) Overview of experimental groups and treatment plan. B) Schematic of the site of whole-cell recording from PFC layer 5 pyramidal neurons and a microscopy image of a patched cell expressing hM3Dq-mCherry (magenta). C-E) Quantitative analysis of membrane properties (resting membrane potential, Vrest; membrane resistance, Rm; membrane capacitance, Cm) showing that intrinsic changes that occur upon prolonged prefrontal hyperactivity (CPA) in PFC pyramidal neurons are prevented or attenuated in the OXTR agonist-treated (RO6958375, RO) group. To test whether these intrinsic changes affect neuronal excitability, current injections were performed. F-I) Representative voltage traces at 0, 20, 40, 340 and 480 pA current injections. J, K) Quantitative analysis of the number of action potentials (APs) produced during each current step and the calculated rheobase (current needed to produce the first AP), reveals that the intrinsic excitability is dramatically reduced under CPA conditions and attenuated in the OXTR agonist-treated group. To probe whether the decreased excitability is associated with a reduced input resistance the current-voltage (IV) relationship was analyzed: L, M) Quantitative analysis of the IV-curve and the calculated input resistance (Rin) at 80 pA, shows that following CPA neurons need more current to produce the same membrane voltage which is associated with a strongly reduced input resistance. Remarkably, this deficit is not detectable in the OXTR agonist-treated group. One-way ANOVA with Tukey’s post-hoc test was performed to test for statistical differences between groups for one condition. Two-way ANOVA with Sidak’s post-hoc test was performed for statistical comparison in J and L, in which two conditions were regarded (statistical differences from post-hoc tests are indicated in the corresponding color for the condition and refer to a difference to control values). In addition, to compare the overall difference between conditions, respective AUCs were tested with one-way ANOVA and Tukey’s post-hoc test (statistical difference is depicted at the side of the legend). * p < 0.05, ** < 0.01, *** p < 0.001. N) Heat map showing gene clusters (annotated using gene sets from: MetaBase, GeneOntology, REACTOME, and the Roche Molecular Phenotyping collection). Every network of category is categorized, for each contrast, by a significance score (positive for up-regulation, negative for down-regulation). As a first pass, for every gene set, we use the maximum of all the absolute scores to sort them. We only include gene sets with at least 3 genes and not more than 500 genes contributing. Based on this, we continue analyses with gene sets scoring at least ±5 in any one of the contrasts analyzed. O) Social behavior data shows a significant decrease in social interaction time with chronic PFC activation, an effect not observed in chronic PFC activation rats treated with OXTR agonists at 0.001 mg/kg or 0.07 mg/kg. Data was analyzed using one-way ANOVA followed by Fisher’s LSD post-hoc test. * p < 0.05. Data in box-and-whiskers plots is displayed as the median with min. to max. All other data is displayed as mean ± SEM (box).

### Sub-chronic selective OXTR activation partially normalizes CPA-induced transcriptomic changes

To better understand the molecular effects of CPA and selective OXTR agonism, we performed transcriptome analysis of tissue we collected following acute slice electrophysiology. RNA-Seq revealed wide-range transcriptomic changes in CPA rats, which were partially prevented with RO6958375 treatment (Fig. 5N). Pathway analysis showed that gene networks associated with mitochondrial function, peptide metabolism, steroid hormones, muscarinic and amine ligand binding receptors, as well as axonal degeneration and toxicity were upregulated following CPA, whereas genes associated with transcriptional regulation of FOXF1 downstream activation were downregulated (Fig. 5N, first column). RO6958375 treatment normalized the expression of a subset of gene networks, involved in mitochondrial function, peptide metabolism, steroid hormones and axonal degeneration (Fig. 5N third column, lower half). However, RO6958375 treatment also augmented expression in certain gene networks, involved in developmental processes, nerve pulse transmission, glutamatergic metabolism and transmission, as well as several neurotransmitter receptor classes (Fig. 5N third column, lower half). Beyond gene network alterations, we observed transcriptional dysregulation for several potassium channels and associated regulatory elements as well as sodium channels, which may contribute to the CPA-induced neuronal hypoexcitability and that were prevented with RO6958375 (Supplementary fig. 3).

### Sub-chronic selective OXTR activation alleviates deficits in social interaction

Finally, we addressed whether the RO6958375-mediated restoration of PFC physiology and transcription also manifest on the behavioral level using a separate group of animals. Social behavior was again evaluated by using the three-chamber sociability test after CPA with RO6958375 or vehicle treatment. Indeed, the social deficits observed in CPA rats were not evident following RO6958375 treatment at 0.001 or 0.07 mg/kg compared to the vehicle treatment (Fig. 5O). Our results demonstrate the prosocial effect of selective OXTR agonism in this circuit model of social behavior impairment, substantiating its translational value.

## Discussion

Modeling social impairment seen in neurodevelopmental and neuropsychiatric conditions preclinically remains a challenge owing to the largely idiopathic nature of these conditions. To address this problem, we adopted a two-step approach, by developing a mechanistic model (the CPA rat model) of social dysfunction and utilizing our novel OXTR-specific agonist to demonstrate efficacy. The CPA model is based on chronic perturbation of the PFC, a candidate node in the social brain network also disrupted in SZ and ASD (see introduction). Previous circuit studies have delineated various neural pathways necessary and sufficient for orchestrating social behavior on acute timescales (2,8). However, the utility of these circuit models in drug discovery remains limited due to the acute nature of these manipulations and the lack of validated pharmacological intervention paradigms. In this context, our work provides a unique systems biology framework where chronic dysfunction of PFC social-circuits is modeled preclinically and can potentially be used to gain insights into the mechanism-of-action of novel therapeutic interventions.

Several lines of evidence suggest that PFC E/I imbalance represents a common circuit substrate for various causes of social impairment in ASD (32,33). Clinical evidence comes from gene mutations affecting excitability and synaptic function in ASD (34). In fact, 50-70% of children with ASD display physiological abnormalities indicative of cortical hyperactivity (35,36). Elegant experimental work has demonstrated that acute PFC hyperactivity is reversibly impeding social interactions in rodents (13,14). We hypothesized that any perturbation of PFC function will also impair social behavior on longer time scales. Therefore, we chose a chemogentic-based approach stimulating PFC activity over several days. Our finding that prolonged hyperactivity of PFC pyramidal neurons results in persistent hypoexcitability has important implications for any experiments involving repeated modulation of neuronal activity via hM3Dq. To our knowledge, this is the first study revealing that hM3Dq-mediated hyperactivity downscales neuronal excitability reminiscent of homeostatic plasticity. In homeostatic plasticity, increased activity levels lead to a reduction of synaptic efficacy or intrinsic excitability to maintain an internal set point, e.g. the firing frequency or membrane potential (reviewed in (37)). In a disease context, similar ‘adaptive’ changes are known from epilepsy, in which network hyperactivity leads to a downscaling of neuronal excitability, due to upregulation of potassium channels (e.g. Kir2.1, HCN and Kv1.1 channels). The resulting increased potassium leakage reduces the resting membrane potential and input resistance, which impairs action potential generation (38–41). We propose that similar mechanisms are engaged upon CPA, in which the hypoexcitability is associated with decreased resting membrane potential, input resistance and elevated potassium currents. Our transcriptome results corroborate this notion, showing that several genes encoding for potassium channels, associated regulatory elements and other ion channels are dysregulated following CPA. While assessing the specific role of individual transcriptional alterations was not in the scope of the present study, and our bulk transcriptomic approach does not delineate cell type-specific effects, our results indicate complex ion channel alterations that may underlie CPA-mediated hypoexcitability. Importantly, gene network alterations involved in mitochondrial function and metabolism may also contribute to hypoexcitability, considering for instance that deficits in energy production will impair neuronal function. While the functional consequences on the network level were not addressed in the current study, it is tempting to speculate that CPA disrupts processing of afferent signals (e.g. social cues) in the PFC, which in turn impedes socially-directed behavior. Notably, our experiments showed that in the LHb and VTA (14,42,43) spontaneous activity was not affected by CPA, which suggests that CPA impairs social behavior predominantly by disrupting PFC physiology, and not necessarily its downstream targets.

Intriguingly, our electrophysiological data demonstrated that repeated OXTR activation prevented or reversed CPA-induced dysfunction of PFC pyramidal neurons, a cell type not known to express OXTRs. A plausible candidate for an indirect mechanism-of-action relies on neuronal inhibition. In line with the concept of homeostatic plasticity, enhanced inhibition could counteract the hyperactivity-induced downscaling of excitability by adjusting the internal set point of affected neurons. Indeed, oxytocin has been shown to boost inhibition in several brain areas, including the PFC (27), the amygdala (25,44), the auditory cortex (45) and other sensory systems (reviewed in (46)). In the PFC, OXTRs are expressed by a subset of GABAergic interneurons (47), that could be activated via OXTR-signaling (reviewed in (16,48)). Indeed, optogenetic activation of these interneurons leads to inhibition of PFC pyramidal neurons (27). We suggest that this inhibition drives the internal set point of hyperactive PFC neurons away from their pathological state to restore PFC function and social behavior.

So far, studying the therapeutic effect of oxytocin has been challenging due to the short half-life, poor brain penetration and the inherent exposure variability associated with administration methods such as intranasal delivery. Therefore, the question remains whether enough oxytocin reaches the brain after peripheral administration to exert behavioral effects (49). Moreover, the interpretation of behavioral effects seen in studies using oxytocin may be hampered due to its agonistic activity on V1aR. To overcome these limitations we developed RO6958375, a derivative of oxytocin that shows outstanding selectivity for OXTR and sustained brain exposure. Our results show that already a low dose of 0.001 mg/kg RO6958375 is sufficient to elicit prosocial effects in our rodent model. In contrast, previous *in vivo* studies (20,50) typically use high oxytocin doses (1 mg/kg i.p.), which could result in OXTR-desensitization and peripheral V1aR activation, causing vasoconstriction that may interfere with behavior readouts. An open question in the field remains as to the right time for oxytocin-treatment in neurodevelopmental disorders, with many advocating for an as-early-as-possible treatment. From our data, indeed, it remains unclear if OXTR activation is only efficacious in a certain time window during which CPA-driven alterations emerge or if treatment could even restore PFC physiology after chronification.

Oxytocin-release in the brain is well known to regulate several aspects of social behavior through a distributed network of OXTR-containing neurons, which receive oxytocinergic input mainly from the paraventricular nucleus (reviewed in (48)). Studies in rodents showed that the PFC, the VTA, the amygdala and the nucleus accumbens are important network nodes to mediate acute prosocial and anxiolytic effects of oxytocin (25,29,51,52). In fact, it has been hypothesized that this may be specifically mediated by oxytocin acting on OXTRs (52,53). While these studies can explain the acute prosocial effects of oxytocin, our study reveals long-lasting restoration of PFC function and transcription. Oxytocin has also been shown to have long-lasting effects on fear and social anxiety (reviewed in (18)), which could be mediated by plasticity-related gene expression downstream of OXTR-signaling (reviewed in (48)). Further investigations into the effect of OXTR-signaling on longer timescales would help to elucidate the therapeutic potential of OXTR agonists for the treatment of behavioral deficits in neurodevelopmental and neuropsychiatric disorders.

In conclusion, our study indicates that hyperactivity-dependent maladaptation could be a critical component in social circuit dysfunction relevant for ASD and SZ (32,54–56), and that selective OXTR activation could prevent the chronification of such deficits.

## Acknowledgements

We would like to acknowledge Michael Weber, Roland Schmucki, Megana Prasad, Michael Saxe, and Audrey Genet for excellent technical assistance. Further, we would like to thank the Roche Internship for Scientific Exchange (RiSE) program for the funding of Philipp Janz.

