## Supplementary material for "Selective oxytocin receptor activation prevents prefrontal circuit dysfunction and social behavioral alterations in response to chronic prefrontal cortex activation in rats"

Authors and affiliations

Philipp Janz

Roche Pharma Research and Early Development, Neuroscience and Rare Diseases Discovery & Translational Area, Roche Innovation Center Basel, F. Hoffmann-La Roche AG, Grenzacherstrasse 124, 4070 Basel

Frederic Knoflach

Roche Pharma Research and Early Development, Neuroscience and Rare Diseases Discovery & Translational Area, Roche Innovation Center Basel, F. Hoffmann-La Roche AG, Grenzacherstrasse 124, 4070 Basel

Konrad Bleicher

Roche Pharma Research and Early Development, Therapeutic Modalities, Roche Innovation Center Basel, F. Hoffmann-La Roche AG, Grenzacherstrasse 124, 4070 Basel

Sara Belli

Roche Pharma Research and Early Development, Pharmaceutical Science, Roche Innovation Center Basel, F. Hoffmann-La Roche AG, Grenzacherstrasse 124, 4070 Basel

Barbara Biemans

Roche Pharma Research and Early Development, Neuroscience and Rare Diseases Discovery & Translational Area, Roche Innovation Center Basel, F. Hoffmann-La Roche AG, Grenzacherstrasse 124, 4070 Basel

Patrick Schnider

Roche Pharma Research and Early Development, Therapeutic Modalities, Roche Innovation Center Basel, F. Hoffmann-La Roche AG, Grenzacherstrasse 124, 4070 Basel

Martin Ebeling

Roche Pharma Research and Early Development, Pharmaceutical Science, Roche Innovation Center Basel, F. Hoffmann-La Roche AG, Grenzacherstrasse 124, 4070 Basel

Christophe Grundschober *

Roche Pharma Research and Early Development, Neuroscience and Rare Diseases Discovery & Translational Area, Roche Innovation Center Basel, F. Hoffmann-La Roche AG, Grenzacherstrasse 124, 4070 Basel

Madhurima Benekareddy *

Roche Pharma Research and Early Development, Neuroscience and Rare Diseases Discovery & Translational Area, Roche Innovation Center Basel, F. Hoffmann-La Roche AG, Grenzacherstrasse 124, 4070 Basel; Calico Life Sciences, 1170 Veterans Blvd., South San Francisco, CA 64080

(* co-last authors)

Corresponding author's name and email address

Name: Philipp Janz

Postal address: Roche Innovation Center Basel, Grenzacherstrasse 124, 4070 Basel, Switzerland

Name: Madhurima Benekareddy

Postal address: Calico Life Science, 1170 Veterans Blvd., South San Francisco, CA 64080, USA

**Extended Material & Methods**

### Viral injections

Rats were deeply anesthetized using a balanced anesthesia protocol (fentanyl 0.005 mg/kg + medetomidine 0.15 mg/kg + midazolam 2 mg/kg). Adeno-associated viruses (AAV) packaged by the Viral Vector Facility of the ETH Zürich were injected stereotaxically into the medial PFC (coordinates relative to bregma: Anterior-posterior, AP = +2.73 mm; medio-lateral, ML = ±0.6 mm; dorso-ventral, DV = -4.5 mm; 900 nL of AAV8-mCaMKIIa-hM3D(Gq)_mCherry-WPRE-hGHp) to express the depolarizing designer receptor exclusively activated by designer drug (DREADD) hM3D. For control rats, a control virus carrying only the genomic sequence for the fluorophore, but not for the DREADD, was used. In a subset of rats a retrograde-infecting AAV was additionally injected in the dorsomedial thalamus (coordinates: AP = +3.20 mm, ML = ±0.65 mm, DV = -5.8 mm; 150 nL of AAV-retro-CAG-EGFP-WPRE-SV40p(A)) to label a subpopulation of PFC projection neurons for subsequent whole-cell recordings.

### Peptide synthesis

RO6958375 was synthesized via microwave-based solid phase Fmoc-chemistry using the Liberty Lite system (CEM, Matthews, US). Synthesis scale was executed at 0.25 mmol with coupling times of 5 minutes per amino acid at elevated temperature (78 °C) applied. The synthesis was carried out using TentalGel-S RAM resin as a solid support (0.24 meq/g). All amino acids used were dissolved in DMF to a 0.2 mol/L concentration. A mixture of HOBT/HBTU 1:1 (0.5 mol/L, 4 eq.) and DIPEA (4 eq.) was used to activate the amino acids. Fmoc-cleavage was achieved with piperidine in DMF (20%) for 3 min. The Fmoc-cleavage step was repeated. For Alloc- and allyl-cleavage, the resin was treated manually with a solution of 20 eq. of phenylsilane and 0.05 eq. of tetrakis(triphenylphosphine)palladium(0) in DCM (5 ml) for 30 min at RT. This procedure was repeated. The resin was washed with a solution of 0.5% sodium dithiocarbamate in DMF twice. The washing step was repeated with DCM. For on-bead cyclisation, the coupling-reagent (4 ml of an 0.5 mol/L solution HOBT/HBTU (1:1) and 1 ml of DIPEA (4 eq.) in DMF was added to the resin. The slurry was shaken for 8 h at RT. The resin was washed with DMF and DCM twice. Completion of cyclisation was verified via ninhydrin test. To cleave the peptide from the resin, 10 ml of the cleavage-cocktail (TFA/TIS/water, ratio of: 95/2.5/2.5) was added to the resin, and the mixture shaken at RT for 1h. Cleaved peptide was precipitated from cold ether (−18 °C). The peptide was centrifuged, and the precipitates were washed twice with cold ether. The precipitate was dissolved in water/acetonitrile and lyophilized. Finally, the crude peptide was purified by preparative HPLC on a Reprospher 100 C18-T Column (100×4.6 mm, Sum particle size). As an eluent system a mixture of 0.1% TFA/water/acetonitrile was used with a gradient of 0-50% acetonitrile within 0-30 min. The fractions were collected and checked by analytical HPLC. Fractions containing pure product were combined and lyophilized. 7.2 mg of white powder was obtained. The product was analyzed via Electrospray Mass Spectrometry with a mass observed of 989.3 (M+H+: expected 989.1).

### Pharmacological characterization of OXTR agonist RO6958375

The peptidic OXTR agonist RO6958375 was profiled in a calcium flux *in vitro* assay on cells expressing human or rat OXTR or the related human or rat vasopressin V1a, V1b and V2 receptors as previously described (32). In a variation to the published protocol, cells were plated for 24 h at 50,000 cells/well in clear-bottomed 96-well plates, washed and dye-loaded for 2 hours with FLIPR calcium 6 no wash assay kit (Molecular Device) in assay buffer (1x Hanks Balanced Salt Solution, 20 mM Hepes) without adding probenicid. The plates were loaded on a fluorometric imaging plate reader (FLIPR) to measure calcium flux after addition of agonist.

**Peptide *in vivo* distribution studies**

Adult female or male Wistar rats weighing circa 250 g were obtained from Harlan Laboratories (Horst, The Netherlands) and housed in a controlled environment (temperature, humidity, and 12-h light/ dark cycle) with access to food and water ad libitum. All rodent studies were conducted with the approval of the local veterinary authority in adherence to the Swiss federal regulations on animal protection and to the rules of the Association for Assessment and Accreditation of Laboratory Animal Care International (AAALAC). Rats were administered RO6958375 subcutaneously at 0.03, 0.1, 0.3, or 4 mg/kg to cover a large range of exposure levels. Serial blood and one CSF sample were collected from each animal under deep anesthesia with 5% isoflurane in pure oxygen. Due to the expected very rapid apparent elimination kinetic, blood sample collection was performed from 0.25 to 4.5 h post-dose, with CSF sampling from 0.8 to 4.5 h. Blood samples were collected by heart puncture into K2EDTA coated polypropylene tubes and placed on ice. Plasma was prepared within 30 min by centrifugation at 3000 g for 5 min at 4 °C and frozen immediately. Between 50 and 100 uL of CSF were collected by cisternal puncture of the atlanto-occipital membrane with a 22-gauge needle through silastic tubing in a 96- well plate. All samples were stored at 20 °C. Compound concentrations in plasma and CSF were determined by means of LC–MS/MS. Merged PK data were analyzed by non-compartmental analysis and PK modeling (details not shown).

### Three-chamber sociability test

Rats were placed in a 3-chamber task from Noldus ethovision (Noldus, Wageningen, NL). At each end of the rectangular box are the enclosures for the social stimulus (unfamiliar conspecific) or the non-social stimulus (unfamiliar object). At the beginning of the test, the rat is placed in the center chamber and allowed to explore the box for 10 mins (habituation). After that, a stimulus rat from a different cage is placed in one of the enclosures and the unfamiliar object in the opposing enclosure. The test rat is allowed to explore the environment for 10 mins (test phase). At the end of 20 minutes, the test rat and stimulus rat were taken out of the arena and the arena was cleaned with 30% alcohol before the next trial. Video was recorded, tracked and analyzed using Ethovision. The light level in the box is maintained at 20 Lux with indirect lighting. Data are expressed as sociability index = 100 x (time in social interaction/total interaction time) - 50. A social interaction is counted when the nose point is in direct proximity to the enclosure with the stimulus rat.

### Electrophysiology

For terminal experiments rats were deeply anesthetized with 2.0% isoflurane (Abbott, Cham, CH) and decapitated. The brain was quickly removed and transferred to the slicing chamber filled with ice-cold N*-*methyl-D-glucamine (NMDG) solution containing (in mM) 110 NMDG, 3 KCl, 1.1 NaH_2_PO_4_, 25 NaHCO_3_, 103.02 HCl, 25 D-glucose, 10 *L*-ascorbic acid, 3 pyruvic acid, 0.5 CaCl_2_ * 2H_2_O and 10 MgCl_2_ * 6H_2_O. The brain was chopped into 350 µm thick frontal sections using a VT1000S vibratome (Leica, Wetzlar, GER). Acute slices were recovered in NMDG solution for 15 min at 35 °C and then transferred either to normal ACSF containing (in mM) 124 NaCl, 2.5 KCl, 1.25 KH_2_PO_4_, 26 NaCO_3_, 10 *D*-glucose, 4 sucrose, 2 MgSO_4_ * 7H_2_O and 2.5 CaCl_2_ * 2H_2_O for extracellular recordings, or to ACSF optimized for whole-cell recordings containing (in mM) 124 NaCl, 2.5 KCl, 1 NaH_2_PO_4_, 25 NaHCO_3_, 20 *D-*glucose, 2 CaCl_2_ * 2H_2_O and 1 MgCl_2_ * 6H_2_O. Both solutions were adjusted to 305-310 mOsm. Patch-clamp whole-cell recordings were performed at RT and a perfusion rate of 1.5 ml/min, whereas spontaneous action potentials were recorded at about 35 °C and a perfusion rate of 3 ml/min.

For testing synaptic transmission within the PFC a bipolar stimulation electrode was located in layer 2/3 and a borosilicate pipette (GC150F-10; Harvard Apparatus, Cambridge, USA) filled with a solution of (in mM) 150 NaCl, 3.5 KCl, 10 HEPES, 10 *D-*glucose, 1.3 MgCl_2_ * 6H_2_O, 2.5 CaCl_2_ * 2H_2_O and a resistance of 2.5-3.5 MΩ was placed in layer 5 to record evoked potentials. The recording electrode was lowered 150 µm into the tissue to obtain maximal responses. Stimulation currents were generated with an STG3000 and MC Stimulus II software (Multi Channel Systems, Reutlingen, Germany). The signal was preamplified with a CV-7B electrode holder (Axon Instruments, Molecular Devices, San Jose, California, US) and again amplified and digitized with a MultiClamp 700B (Axon Instruments) using 100xAC membrane potential (200 mV/mV) mode, a bessel filter at 2.4 kHz and alternating current at 1 Hz. Signals were recorded with Clampex software (Axon Instruments). First, an input-output protocol was performed by increasing the stimulation intensity till 300 µA in 20 µA current steps. For electrical pulses we used a square shape and a duration of 100 µs. Then, we performed a paired pulse protocol using intervals of 20, 50, 100, 200, 400 and 800 ms and a stimulation current (typically 80-100 µA) which elicited the half-maximum slope of the postsynaptic field potential (fPSP). Finally, synaptic short term depression (STD) was tested by delivering 40 pulses at 5, 10, 20 and 40 Hz. All protocols were performed in 3 slices for each rat and repeated three times. The amplitudes and slopes of the presynaptic fiber volley (FV) and the fPSP were semi-automatically analyzed using Clampex and averaged per animal.

Spontaneous activity was recorded in the central LHb and the VTA using the same pipette solution and amplifier settings as mentioned above. Active neurons were identified by slowly protruding the electrode forward into the tissue. As soon as recurrent spikes occurred in the voltage trace the electrode position was fine-adjusted to obtain a signal amplitude between 0.2 and 0.5 mV. After waiting 5 min for the spike rate to normalize, action potential firing was recorded for at least 10 min. The firing frequency was analyzed for each neuron by using an adjustable threshold for spike detection in Clampex. Spike shapes were checked for each recording to ensure that the calculated firing frequencies relate to individual neurons.

For patch-clamp experiments PFC layer 5 pyramidal neurons were morphologically identified and recorded in whole-cell mode. The patch pipette (GC150F-10; Harvard Apparatus) contained a solution of (in mM) 145 KMeSO_4_, 10 Hepes, 10 NaCl, 10 EGTA, 5 MgATP, 0.5 Na_2_ATP, 1 CaCl_2_ * 2H_2_O and 1 MgCl_2_ * 6H_2_O (adjusted to 290 mOsm and pH 7.2) and had a resistance 2.5-3.5 MΩ. Using positive pressure neurons were visually approached. By applying negative pressure a gigaseal was formed with the targeted neuron. Then, the cell membrane was opened and protocols were started 10 min after opening, to allow for sufficient exchange of intracellular solution and normalization of cell physiology. Cell viability was tested before and after each trial. The membrane potential was held at -70 mV, and only neurons for which leak currents did not exceed ±100 pA and for which the axial resistance did not show changes higher than 20% were included for analysis. After inferring the basic neuronal properties (i.e. the membrane resistance, R_m_; the membrane capacity, C_m_; and the resting membrane potential, V_rest_), steady-state potassium currents were assessed by holding the membrane potential at -40 or 0 mV for 500 ms respectively. Intrinsic excitability was inferred by quantifying the number of evoked generated action potentials during incremental current steps of 20 pA (100 ms duration) until a stimulus intensity of 500 pA was reached.

### RNA Sequencing

The mPFC was dissected out from the sections after electrophysiology, and the tissue was flash frozen in liquid nitrogen and stored at -80 °C until further use. RNA was extracted using Qiagen RNeasy mini kit (Qiagen, Hilden, GER). RNA quality was checked using Agilent RNA 6000 Nano Kit (Agilent Technologies, Santa Clara, US), RNA concentration was measured with Nanodrop (Thermo Fisher Scientific, Waltham, US). 500 ng of RNA was used as input for library preparation using the Illumina TruSeq Stranded mRNA library prep kit (Illumina, San Diego, US) according to the manufacturer's instructions. Libraries were quality checked on a Fragment Analyzer (Agilent Technologies) and concentration was measured with Qubit dsDNA HS Assay kit (Thermo Fisher Scientific). Libraries were pooled and sequenced on a HiSeq4000 (Illumina).

**Supplementary Figures & Tables**

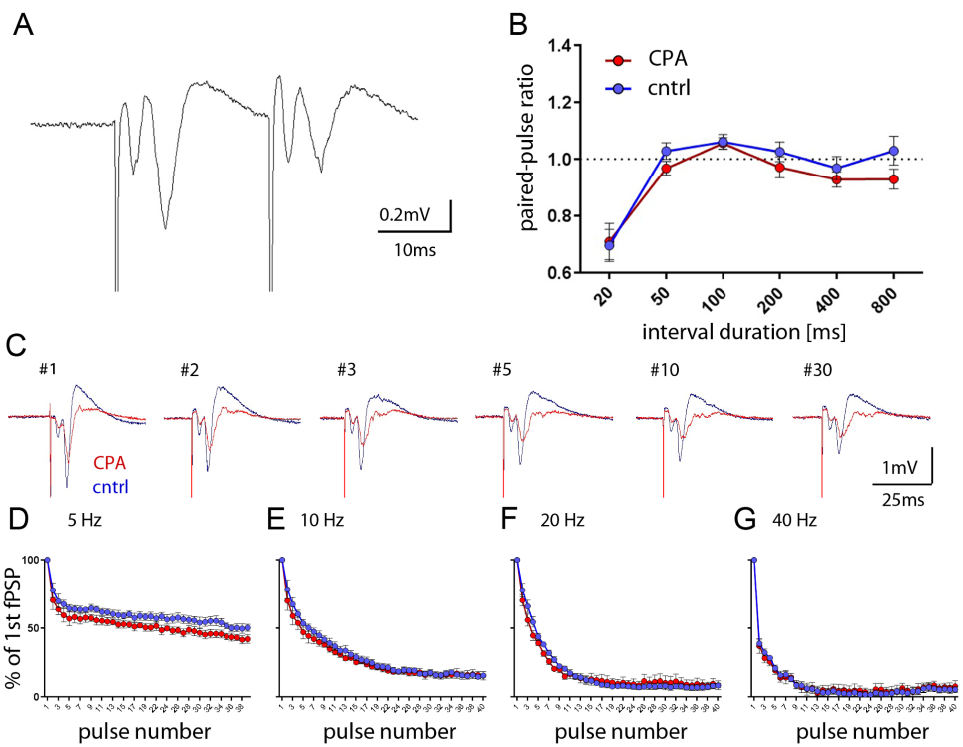

**Supplementary figure 1: Pre-pulse responses and short-term plasticity are preserved in PFC.** A) Representative voltage trace of a paired-pulse response at 20 ms inter-stimulus-interval duration. B) Quantitative analysis of paired-pulse ratios show no significant differences between groups. C) Representative evoked responses from a short-term depression protocol (here with 40 pulses presented at 5 Hz). D-G) Synaptic depression was quantified by inferring the %-change of the respective fPSP amplitude to the first fPSP. No significant differences are evident between groups for different stimulus frequencies (5, 10, 20 and 40 Hz). For paired-pulse and short-term depression protocols the stimulation intensity was used that elicited half maximum slope of the response (80-100 uA). Two-way ANOVA with Sidak`s post-hoc test was performed for statistical comparison. Data is displayed as mean ± SEM.

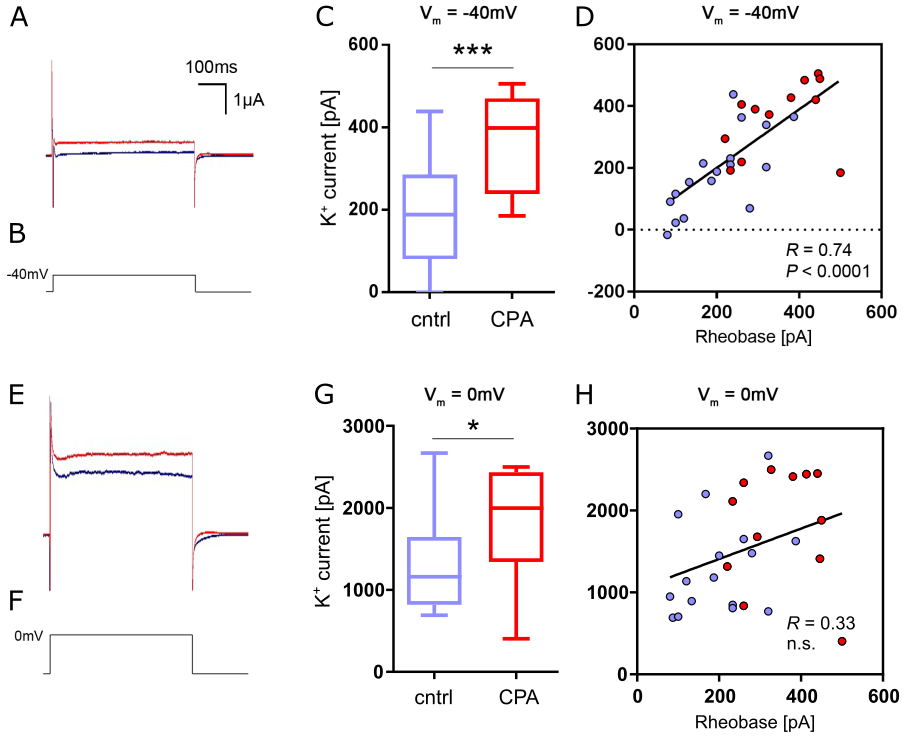

**Supplementary figure 2: Increased potassium currents.** A, B) Representative current traces at -40 mV membrane potential. Steady- state current represents the potassium current mediated by a subset of voltage-gated potassium channels. C) Quantitative analysis of the potassium steady-state current shows a highly significant increase for the CPA group. D) Linear regression analysis shows that this increase is positively correlated with the neuronal excitability represented by the rheobase. E-H) Same analysis performed for membrane potentials at 0 mV. Overall group comparison was performed with Student`s t-test. * p < 0.05, ** < 0.01, *** < 0.001. Linear regression analysis was done with Spearman’s correlation. Data is displayed as the median with min. to max. (box-and-whiskers).

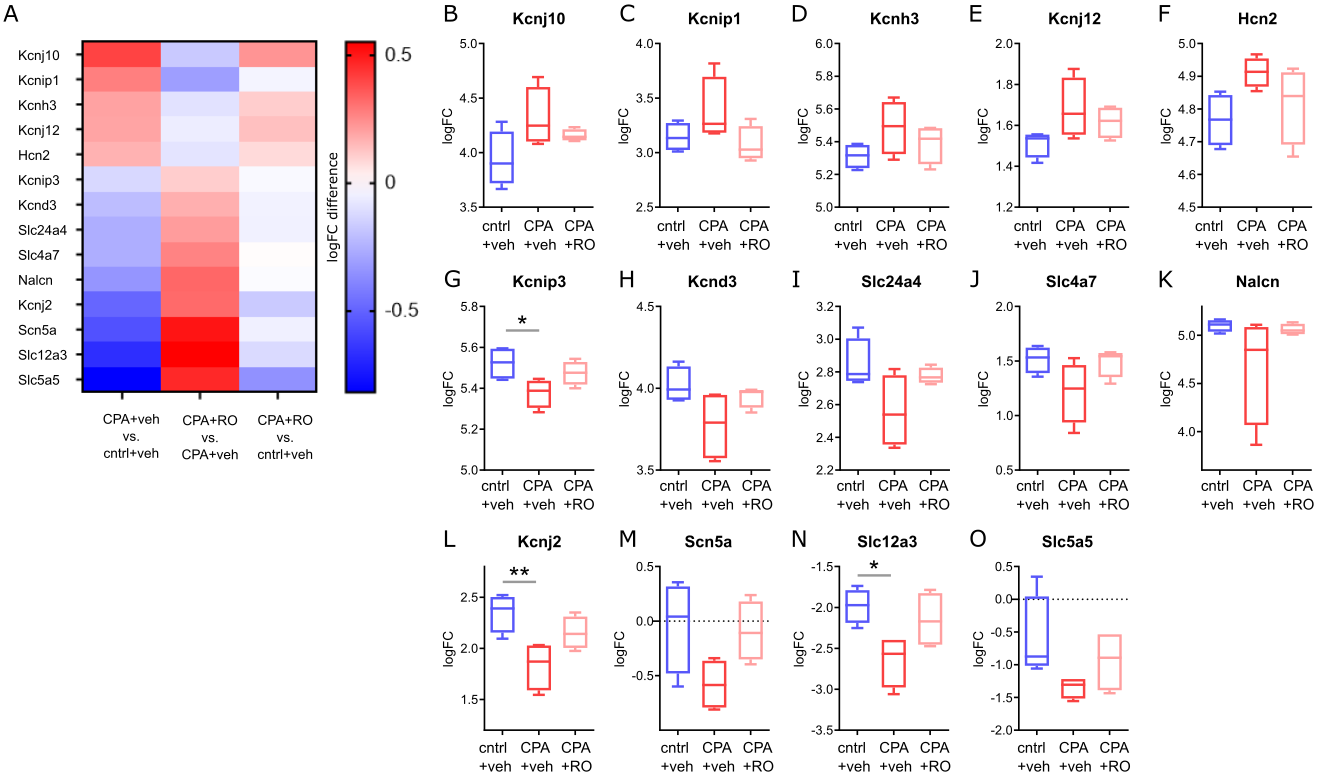

**Supplementary figure 3: Transcriptomic alterations focusing on ion channels involved in regulating cellular excitability.** A) Heatmap displaying significantly altered expression (between CPA+veh and control+veh, first column) of ion channels involved in cellular excitability. Countering of CPA-induced differential gene expression is evident followed by OXTR agonist (RO6958375, RO) treatment (second column), resulting in a normalization compared to control (third column).  B) Box plots for individual genes represented in the heatmap showing the fold change of gene expression in different conditions. Tested with one-way ANOVA and Tukey`s post-hoc test. * p < 0.05, ** < 0.01. Data is displayed as mean ± SEM.

**Supplementary table 1: Numeric values and statistical tests for key results**.

| **Figure** | **Comparison** | **Mean 1 ± SEM** | **Mean 2 ± SEM** | **p value** | **statistical test** |
| --- | --- | --- | --- | --- | --- |
| Fig. 1C | hM3Dq baseline vs. hM3Dq+CNO | 21.5 ± 4.1  (8 rats) | 9.6 ± 3.2  (8 rats) | 0.066 | 2-way ANOVA Fisher’s LSD |
|  | control+CNO vs. hM3Dq+CNO | 24.6 ± 4.5  (9 rats) | 9.6 ± 3.2  (8 rats) | < 0.05 | 2-way ANOVA Sidak’s test |
| Fig. 1H | AUC of control vs. AUC of CPA | 13.24 ± 0.82 mV  (4 rats) | 9.84 ± 0.65 mV  (5 rats) | < 0.05 | Unpaired Student’s t-test |
| Fig. 2B | control vs. CPA | -61.62 ± 1.97 mV  (21 cells, 4 rats) | -67.82 ± 1.89 mV  (17 cells, 4 rats) | < 0.05 | Unpaired Student’s t-test |
| Fig. 2C | control vs. CPA | 83.1 ± 6.36 MΩ  (21 cells, 4 rats) | 61.29 ± 6.89 MΩ  (17 cells, 4 rats) | < 0.05 | Unpaired Student’s t-test |
| Fig. 2H | AUC of control vs. AUC of CPA | 90.76 ± 18.48  (17 cells, 4 rats) | 17.24 ± 5.56  (12 cells, 4 rats) | < 0.01 | Unpaired Student’s t-test |
| Fig. 2I | control vs. CPA | 202.8 ± 22.4 mV  (17 cells, 4 rats) | 351.8 ± 28.2  (12 cells, 4 rats) | < 0.001 | Unpaired Student’s t-test |
| Fig. 2M | AUC of control vs AUC of CPA | 140 ± 15.41  (15 cells, 4 rats) | 75.14 ± 8.01  (12 cells, 4 rats) | < 0.01 | Unpaired Student’s t-test |
| Fig. 2N | control vs. CPA | 177 ± 24.55 MΩ  (16 cells, 4 rats) | 85.14 ± 8.5 MΩ  (12 cells, 4 rats) | < 0.01 | Unpaired Student’s t-test |
| Suppl. Fig. 2C | control vs. CPA | 187.6 ± 31.7 pA  (17 cells, 4 rats) | 365.8 ± 33.4 pA  (12 cells, 4 rats) | < 0.001 | Unpaired Student’s t-test |
| Suppl. Fig. 2G | control vs. CPA | 1313 ± 142.3 pA  (16 cells, 4 rats) | 1815 ± 201.7 pA  (12 cells, 4 rats) | < 0.05 | Unpaired Student’s t-test |
| Fig. 3D | control vs. CPA | 5.79 ± 1.43 Hz  (17 cells, 4 rats) | 9.90 ± 2.80 Hz  (14 cells, 4 rats) | 0.179 | Unpaired Student’s t-test |
| Fig. 3J | control vs. CPA | 1.21 ± 0.15 Hz  (19 cells, 4 rats) | 1.27 ± 0.15 Hz  (13 cells, 4 rats) | 0.784 | Unpaired Student’s t-test |
| Fig. 3K | control vs. CPA | 6.11 ± 1.1 Hz  (8 cells, 4 rats) | 4.58 ± 0.48 Hz  (5 cells, 4 rats) | 0.316 | Unpaired Student’s t-test |
| Fig. 5C | control+veh vs. CPA+veh | -63.66 ± 1.72 mV  (29 cells, 4 rats) | -72.17 ± 1.65 mV  (24 cells) | < 0.01 | 1-way ANOVA Tukey’s test |
|  | control+veh vs. CPA+RO | -63.66 ± 1.72 mV  (29 cells) | -66.87 ± 1.74 mV  (32 cells, 4 rats) | 0.364 | 1-way ANOVA Tukey’s test |
|  | CPA+veh vs. CPA+RO | -72.17 ± 1.65 mV  (24 cells, 4 rats) | -66.87 ± 1.74 mV  (32 cells, 4 rats) | 0.089 | 1-way ANOVA Tukey’s test |
| Fig. 5D | control+veh vs. CPA+veh | -82.38 ± 6.25 MΩ  (32 cells, 4 rats) | 63.6 ± 3.88 MΩ  (25 cells, 4 rats) | < 0.05 | 1-way ANOVA Tukey’s test |
|  | control+veh vs. CPA+RO | -82.38 ± 6.25 MΩ  (32 cells, 4 rats) | 77.59 ± 4.64 MΩ  (37 cells, 4 rats) | 0.774 | 1-way ANOVA Tukey’s test |
|  | CPA+veh vs. CPA+RO | 63.6 ± 3.88 MΩ  (25 cells, 4 rats) | 77.59 ± 4.64 MΩ  (37 cells, 4 rats) | 0.155 | 1-way ANOVA Tukey’s test |
| Fig. 5E | control+veh vs. CPA+veh | 61.31 ± 3.69 pF  (32 cells, 4 rats) | 79.54 ± 4.71 pF  (26 cells, 4 rats) | < 0.01 | 1-way ANOVA Tukey’s test |
|  | control+veh vs. CPA+RO | 61.31 ± 3.69 pF  (32 cells, 4 rats) | 65.11 ± 3.48 pF  (36 cells, 4 rats) | 0.754 | 1-way ANOVA Tukey’s test |
|  | CPA+veh vs. CPA+RO | 79.54 ± 4.71 pF  (26 cells, 4 rats) | 65.11 ± 3.48 pF  (36 cells, 4 rats) | < 0.05 | 1-way ANOVA Tukey’s test |
| Fig. 5K | control+veh vs. CPA+veh | 168.8 ± 20.59 mV  (25 cells, 4 rats) | 329.6 ± 22.56 mV  (23 cells, 4 rats) | < 0.001 | 1-way ANOVA Tukey’s test |
|  | control+veh vs. CPA+RO | 168.8 ± 20.59 mV  (25 cells, 4 rats) | 209 ± 22.38 mV  (29 cells, 4 rats) | 0.388 | 1-way ANOVA Tukey’s test |
|  | CPA+veh vs. CPA+RO | 329.6 ± 22.56 mV  (23 cells, 4 rats) | 209 ± 22.38 mV  (29 cells, 4 rats) | < 0.001 | 1-way ANOVA Tukey’s test |
| Fig. 5J | AUC of control+veh vs. AUC of CPA+veh | 16.56 ± 4.11  (25 cells, 4 rats) | 3.09 ± 1.65  (23 cells, 4 rats) | < 0.01 | 1-way ANOVA Tukey’s test |
|  | AUC of control+veh vs. AUC of CPA+RO | 16.56 ± 4.11  (25 cells, 4 rats) | 9.79 ± 2.74  (29 cells, 4 rats) | 0.249 | 1-way ANOVA Tukey’s test |
|  | AUC of CPA+veh vs. AUC of CPA+RO | 3.09 ± 1.65  (23 cells, 4 rats) | 9.79 ± 2.74  (29 cells, 4 rats) | 0.271 | 1-way ANOVA Tukey’s test |
| Fig. 5L | AUC of control+veh vs. AUC of CPA+veh | 38.64 ± 3.69  (29 cells, 4 rats) | 24.37 ± 1.89  (24 cells, 4 rats) | < 0.05 | 1-way ANOVA Tukey’s test |
|  | AUC of control+veh vs. AUC of CPA+RO | 38.64 ± 3.69  (29 cells, 4 rats) | 36.9 ± 4.56  (32 cells, 4 rats) | 0.940 | 1-way ANOVA Tukey’s test |
|  | AUC of CPA+veh vs. AUC of CPA+RO | 24.37 ± 1.89  (24 cells, 4 rats) | 36.9 ± 4.56  (32 cells, 4 rats) | 0.063 | 1-way ANOVA Tukey’s test |
| Fig. 5M | control+veh vs. CPA+veh | 198.6 ± 20.1 MΩ  (29 cells, 4 rats) | 119.8 ± 9.4 MΩ  (24 cells, 4 rats) | < 0.05 | 1-way ANOVA Tukey’s test |
|  | control+veh vs. CPA+RO | 198.6 ± 20.1 MΩ  (29 cells, 4 rats) | 180.7 ± 20.5 MΩ  (32 cells, 4 rats) | 0.758 | 1-way ANOVA Tukey’s test |
|  | CPA+veh vs. CPA+RO | 119.8 ± 9.4 MΩ  (24 cells, 4 rats) | 180.7 ± 20.5 MΩ  (32 cells, 4 rats) | 0.063 | 1-way ANOVA Tukey’s test |
| Fig. 5O | control+veh vs. CPA+veh | 188.14 ± 24.6 s  (8 rats) | 119.6 ± 13.3 s  (8 rats) | < 0.05 | 1-way ANOVA Fisher’s LSD |
|  | CPA+veh vs. CPA+RO 0.001mg/kg | 119.6 ± 13.3 s  (8 rats) | 198.6 ± 15.7 s  (8 rats) | < 0.05 | 1-way ANOVA Fisher’s LSD |
|  | CPA+veh vs. CPA+RO 0.07mg/kg | 119.6 ± 13.3 s  (8 rats) | 190.2 ± 23.6 s  (8 rats) | < 0.05 | 1-way ANOVA Fisher’s LSD |

For each figure referenced in the results section mean 1 and mean 2 with standard error of the means (SEM) of the corresponding comparison is provided, including the p values and applied statistical test.

**Supplementary table 2: EC50 values for selectivity tests**.

| **Receptor** | **Compound** | **EC50** |
| --- | --- | --- |
| rat OXTR | RO6958375 | 0.011 ± 0.004 nM (8 cells) |
|  | Oxytocin | 0.0048 ± 0.002 nM (7 cells) |
| rat V1aR | RO6958375 | inactive (7 cells) |
|  | Oxytocin | 4.5 ± 0.8 nM (3 cells) |
|  | Vasopressin | 0.011 ± 0.3 nM (3 cells) |
| human OXTR | RO6958375 | 0.025 ± 0.004 nM (7 cells) |

EC50 values are provided for RO6958375, oxytocin and vasopressin on either OXTR or V1aR showing that RO6958375 strongly activates OXTR but is inactive on V1aR.

**Supplementary table 3: Selectivity testing of the peptidic OXTR agonist RO6958375**

| **Receptor / channels transporter binding assays** | **% Inhibition of Control Specific Binding** | **1st / % of Control Specific Binding** | **2nd / % of Control Specific Binding** | **Mean / % of Control Specific Binding** | **Reference Compound** | **IC50 Ref [M]** | **Ki Ref [M]** |
| --- | --- | --- | --- | --- | --- | --- | --- |
| A1 | **-4** | 109.1 | 99.1 | 104.1 | CPA | 1.80E-09 | 7.20E-10 |
| A3 | **4** | 88 | 104.9 | 96.4 | IB-MECA | 2.70E-10 | 1.60E-10 |
| alpha 1A | **1** | 100.9 | 96.7 | 98.8 | WB 4101 | 2.90E-10 | 1.50E-10 |
| alpha 2A | **-7** | 107.6 | 105.8 | 106.7 | yohimbine | 4.70E-09 | 2.10E-09 |
| beta 1 | **-5** | 106.9 | 103.4 | 105.2 | atenolol | 2.50E-07 | 1.40E-07 |
| AT1 | **-1** | 96.1 | 106.6 | 101.4 | saralasin | 4.00E-10 | 2.00E-10 |
| BZD | **-16** | 115.9 | 116.2 | 116 | diazepam | 8.50E-09 | 7.10E-09 |
| D1 | **-2** | 117.6 | 86.8 | 102.2 | SCH 23390 | 4.20E-10 | 1.70E-10 |
| D2S | **6** | 94 | 93.4 | 93.7 | 7-OH-DPAT | 4.90E-09 | 2.00E-09 |
| glycine | **15** | 85.2 | 84.1 | 84.6 | glycine | 3.20E-07 | 2.90E-07 |
| H1 | **-1** | 102.3 | 99.5 | 100.9 | pyrilamine | 2.20E-09 | 1.40E-09 |
| H2 | **18** | 76.9 | 87 | 81.9 | cimetidine | 5.60E-07 | 5.40E-07 |
| H3 | **-6** | 112.6 | 99.3 | 105.9 | alpha-Me-histamine | 2.70E-09 | 6.60E-10 |
| I1 | **-6** | 94.4 | 116.7 | 105.5 | rilmenidine | 2.70E-07 | 1.40E-07 |
| M2 | **-14** | 119.6 | 108 | 113.8 | methoctramine | 6.90E-08 | 4.80E-08 |
| M4 | **-17** | 116.8 | 117 | 116.9 | 4-DAMP | 4.40E-10 | 2.70E-10 |
| N muscle-type | **5** | 93 | 96.3 | 94.7 | Alpha-bungarotoxin | 2.00E-09 | 1.80E-09 |
| kappa | **-13** | 115.4 | 110 | 112.7 | U 50488 | 1.20E-09 | 8.00E-10 |
| mu | **2** | 101.2 | 95.2 | 98.2 | DAMGO | 1.80E-09 | 7.40E-10 |
| PPARgamma | **-3** | 99.2 | 107 | 103.1 | rosiglitazone | 1.10E-08 | 5.90E-09 |
| PCP | **-6** | 116.2 | 94.8 | 105.5 | MK 801 | 4.70E-09 | 2.60E-09 |
| FP | **3** | 92.2 | 101.7 | 96.9 | PGF2alpha | 3.10E-09 | 2.00E-09 |
| 5-HT1A | **16** | 78.8 | 90 | 84.4 | 8-OH-DPAT | 8.60E-10 | 5.40E-10 |
| 5-HT2A | **9** | 88.6 | 92.6 | 90.6 | (±)DOI | 3.20E-10 | 2.40E-10 |
| 5-HT2B | **-2** | 103.2 | 100.7 | 102 | (±)DOI | 1.40E-08 | 7.00E-09 |
| 5-HT3 | **20** | 78.6 | 81.5 | 80 | MDL 72222 | 5.70E-09 | 4.00E-09 |
| sigma | **4** | 109.9 | 82.6 | 96.2 | haloperidol | 5.90E-08 | 4.80E-08 |
| sst4 | **17** | 85.8 | 79.9 | 82.8 | somatostatin-14 | 1.60E-09 | 1.60E-09 |
| GR | **-9** | 109.3 | 109 | 109.1 | dexamethasone | 2.90E-09 | 1.40E-09 |
| ERalpha | **-2** | 109.6 | 93.9 | 101.7 | 17-beta-estradiol | 5.30E-09 | 4.30E-09 |
| Ca2+ channel | **-36** | 129 | 143 | 136 | diltiazem | 5.40E-08 | 4.20E-08 |
| Na+ channel | **10** | 86.4 | 92.9 | 89.6 | veratridine | 1.10E-05 | 1.00E-05 |
| NE transporter | **3** | 98 | 96.4 | 97.2 | protriptyline | 2.60E-09 | 1.90E-09 |
| 5-HT transporter | **-4** | 106.7 | 100.3 | 103.5 | imipramine | 2.50E-09 | 1.20E-09 |
| **Enzyme functional assays** |  |  |  |  |  |  |  |
| COX2 | **15** | 87.2 | 83.3 | 85.3 | NS 398 | 5.80E-08 | 1.6 |
| PDE5 | **-21** | 130.8 | 112 | 121.4 | dipyridamole | 1.50E-06 | 1.4 |
| ACE | **-28** | 129.2 | 127.2 | 128.2 | captopril | 4.40E-10 | 1.6 |
| HIV-1 protease | **5** | 104.6 | 86.1 | 95.3 | pepstatin A | 1.70E-06 | 2.1 |
| CDK2 | **13** | 98.3 | 75.8 | 87.1 | staurosporine | 1.00E-08 | 1.1 |
| GSK3alpha | **-5** | 104.6 | 105.9 | 105.2 | staurosporine | 3.40E-08 | 1.8 |
| GSK3beta | **-2** | 106.3 | 97.8 | 102.1 | staurosporine | 5.20E-08 | 2 |
| acetylcholinesterase | **-2** | 101.1 | 101.9 | 101.5 | neostigmine | 4.30E-08 | 1.2 |
| MAO-A | **-15** | 123.1 | 107 | 115 | clorgyline | 3.90E-08 | 0.8 |
| MAO-B enzyme | **-14** | 114.1 | 113.6 | 113.8 | deprenyl | 3.30E-08 | 1.4 |
| xanthine oxidase / superoxide O2-scavenging | **8** | 96.1 | 88.2 | 92.1 | allopurinol | 5.60E-06 | 0.9 |

**Supplementary table 3: Selectivity testing of the peptidic OXTR agonist RO6958375**. Testing was performed using radioligand binding and enzyme functional assays on 45 human receptors, channels and enzymes. Results showing an inhibition (or stimulation for assays run in basal conditions) higher than 50% are considered to represent significant effects of the test compounds. 50% is the most common cut-off value for further investigation Results showing an inhibition (or stimulation) lower than 20% compared to control values are not considered significant and mostly attributable to variability of the signal around the control level. Low to moderate negative values have no real meaning and are attributable to variability of the signal around the control level.
